# Duplicated *shh* genes in *Xenopus laevis* reveal an essential role in early limb development

**DOI:** 10.64898/2026.05.25.726107

**Authors:** Norie Kagawa, Rima Mizuno, Yoshihiko Umesono, Ken-ichi T Suzuki, Makoto Mochii

**Affiliations:** Department of Life Science, Graduate School of Science, University of Hyogo, Ako-gun, Hyogo, 678-1297, Japan; Department of Basic Biology, The Graduate University for Advanced Studies, SOKENDAI, Okazaki, Aichi, 444-8585, Japan; Emerging Model Organisms Facility, Trans-scale Biology Center, National Institute for Basic Biology, National Institutes of Natural Sciences (NIBB), Okazaki, Aichi, 444-8585, Japan

**Keywords:** Sonic hedgehog, *Xenopus laevis*, Limb outgrowth, Zone of polarizing activity

## Abstract

Sonic Hedgehog (SHH) is a key signaling molecule controlling anterior-posterior patterning in vertebrate paired appendages, but its requirement for initial appendage outgrowth varies across lineages. Whether SHH is genetically required for early limb-bud outgrowth in amphibians has remained unresolved due to embryonic pleiotropy caused by systemic SHH disruption. Here, we exploited the duplicated homeologs (*shh.L* and *shh.S*) in the allotetraploid frog *Xenopus laevis* to bypass systemic pleiotropy and investigate the limb-specific function of *shh*. We found that the conserved limb enhancer MFCS1/ZRS is retained at the *shh.L* locus but degraded at *shh.S*, resulting in limb-bud expression strictly confined to *shh.L*. Individual CRISPR/Cas9 disruption of *shh.L*, but not *shh.S*, led to a complete failure of initial limb-bud outgrowth and the absence of appendicular skeletal elements, without causing severe systemic defects. Our findings provide direct genetic evidence that *shh.L* is essential for initial limb outgrowth in *X. laevis*, demonstrating that dependence on SHH for early appendage initiation is strongly conserved in teleosts and amphibians but was reduced during amniote evolution.

## Introduction

Sonic Hedgehog (SHH) is a conserved signal controlling anterior-posterior patterning in vertebrate paired appendages, but its requirement during early outgrowth varies across lineages^1^. In mouse *Shh*-null embryos, all four limb buds form and produce proximal skeletal structures despite severe distal defects, demonstrating that initial limb-bud outgrowth can proceed without SHH^2^. Conversely, in teleosts such as zebrafish and medaka, disruption of *shh* coding regions or limb enhancers severely impairs fin-bud outgrowth^3,4,5^. These findings raise the possibility that early limb outgrowth became less dependent on SHH during amniote evolution. Amphibians provide an important comparative perspective on the water-to-land transition. In axolotl, CRISPR disruption of *Shh* prevents forelimb-bud outgrowth, although severe systemic abnormalities make it less straightforward to assess its limb-specific role^6^. Pharmacological inhibition of Hedgehog signaling in *Xenopus tropicalis* can cause complete limb loss^7^. However, whether *shh* itself is genetically required for initial limb-bud outgrowth has remained unresolved. Because anuran limb buds develop post-embryonically, pleiotropic effects of systemic *shh* disruption can obscure its role specifically in early limb outgrowth. In the allotetraploid genome of Xenopus laevis, shh is retained as two homeologs, *shh.L and shh.S*^8^. We exploited the duplicated homeologs to bypass systemic pleiotropy and investigate their respective roles in limb-bud outgrowth.

## Results and discussion

Sequence alignment revealed that the conserved limb enhancer MFCS1/ZRS is present at the *shh.L* locus, whereas the corresponding *shh.S* region exhibits extensive sequence loss encompassing the core enhancer (Figure S1A)^9,10^. Knock-in reporter *X. laevis* (*shh.L:egfp* and *shh.S:egfp*) showed that while both homeologs are active in axial tissues during embryogenesis (Figure S1C and S1D), EGFP expression in developing limb buds is strictly confined to *shh.L:egfp* (Figures 1A, 1B and S1E). Quantitative RT-PCR confirmed robust *shh.L* expression in wild-type limb buds, whereas *shh.S* transcripts were undetectable (Figure S1F). These results demonstrate a clear partitioning of limb expression between the two homeologs, with shh.L acting as the sole homeolog expressed in the limb bud.

**Figure 1.**
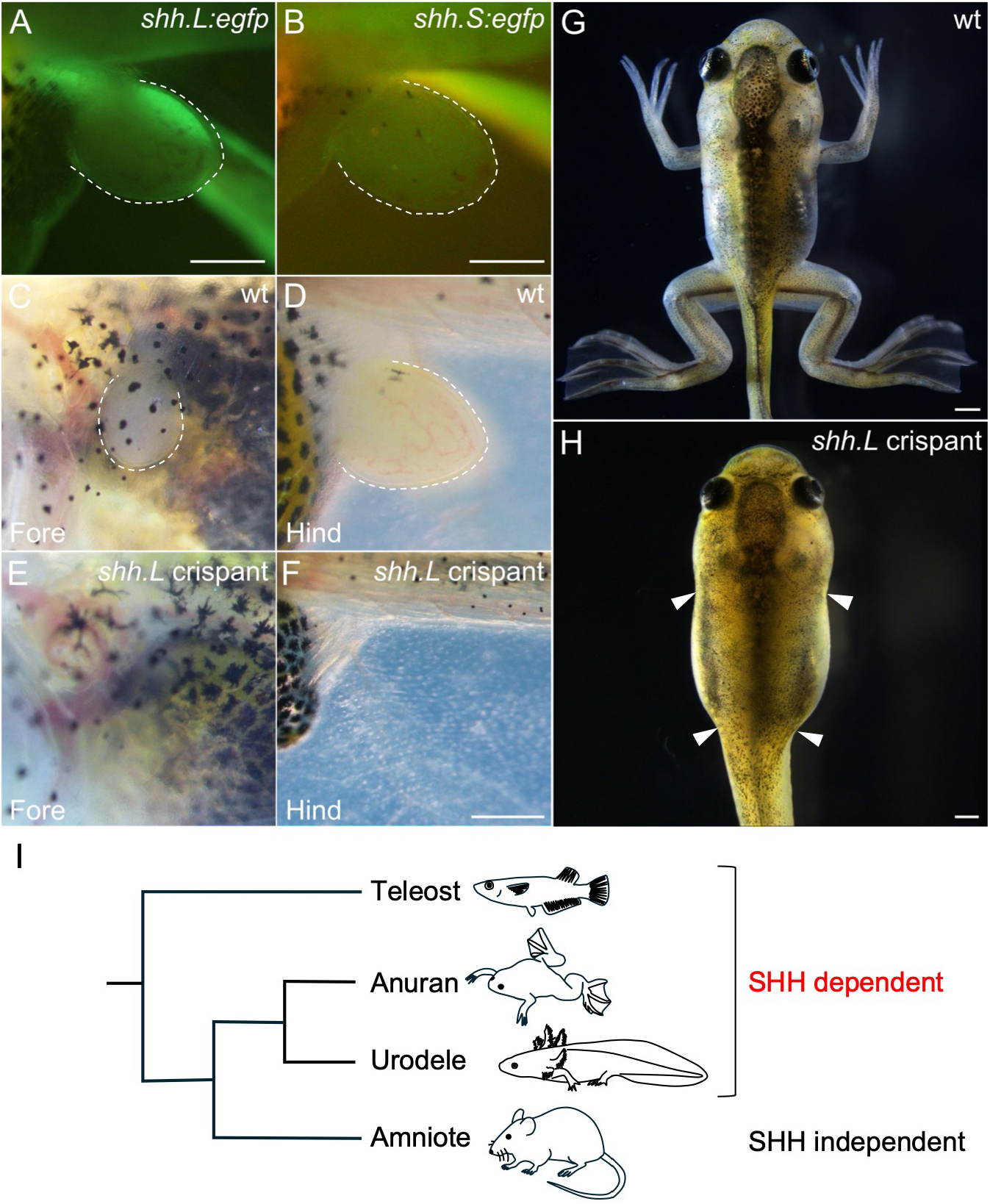
Selective limb-bud expression of *shh.L* and complete limb loss upon its targeted disruption. (A and B) Representative fluorescence images of NF 50 hindlimb buds from *shh.L:egfp* (A) and *shh.S:egfp* (B) tadpoles. Dotted lines indicate the outline of the limb bud. a and p indicate anterior and posterior, respectively. Scale bar: 300 µm. (C–F) Representative images of the forelimb (C and E) and hindlimb (D and F) regions in NF 50 wild-type (C and D) and body-length-matched *shh.L* crispant (E and F) tadpoles. Scale bar: 300 µm. (G and H) Wild-type (G) and *shh.L* crispant (H) tadpoles at the climax stage. Arrowheads indicate the expected positions of the forelimbs and hindlimbs in the *shh.L* crispant, highlighting their complete absence. Scale bar: 1 mm. (I) Schematic illustrating evolutionary changes in SHH dependence during early paired-appendage outgrowth across vertebrates.

Taking advantage of this unique expression profile to bypass systemic pleiotropy, we targeted each homeolog individually using CRISPR/Cas9. Both *shh.L* and *shh.S* crispants completed embryogenesis and developed into viable, feeding tadpoles. All wild-type siblings and *shh.S* crispants formed limb buds and developed to adulthood with normal limbs (n = 13 and n = 15, respectively). By contrast, 9 of 10 *shh.L* crispants lacked one or more limb buds (Figures 1C–1F). Consistent with F_0_ mosaicism (Figure S2B), phenotypes ranged from loss of a single limb to absence of all four limbs (Figures 1G, 1H, S2C and S2D; Supplementary table 3). Serial time-course observations revealed no transient limb-bud outgrowth in crispants exhibiting complete limb loss (n = 4; Figure S2F). These *shh.L* crispants survived to metamorphic climax but remained externally limbless (Figures 1H, S2C and S2D). Skeletal staining confirmed the absence of the corresponding appendicular elements including pelvic girdle in crispants with complete hindlimb loss (n = 5; Figure S2H, Supplementary table 3).

Although we cannot exclude an earlier requirement in limb-field specification, these results demonstrate that *shh.L*, but not *shh.S*, is required for initial limb-bud outgrowth in *X. laevis*. The partitioned expression of the two homeologs allows the limb requirement for SHH to be examined without overt systemic abnormalities that would otherwise confound the analysis. Comparative genetic studies indicate that the requirement for SHH in early paired-appendage outgrowth is not uniform across vertebrates. In zebrafish and medaka, *shh* genes are required for early fin-bud outgrowth^3,4,5^. This requirement for early limb outgrowth is also consistent with findings in axolotls^6^. In sharp contrast, mouse limb buds still form in the absence of SHH^2^. We therefore propose that the dependence of early paired-appendage outgrowth on SHH has been evolutionarily reconfigured, with a strong requirement in teleosts and amphibians but reduced dependence in amniote limbs (Figure 1I).

## Materials and methods

### Animals

Wild-type *X. laevis* adults were obtained commercially. Fertilized eggs were obtained by artificial insemination and staged according to Nieuwkoop and Faber (NF) stage^S1^. All animal care and experimental procedures were approved by the Institutional Animal Care and Use Committee of the University of Hyogo and the National Institutes of Natural Sciences.

### Comparative genomics

The mouse MFCS1/ZRS sequence was downloaded from the NCBI database. To identify homologous regions in *Xenopus*, BLAST searches were performed on the Xenbase platform (https://www.xenbase.org). Interspecific sequence comparisons were conducted using VISTA tools (https://genome.lbl.gov/vista/index.shtml).

### Plasmid DNA construction, Cas9 protein, and sgRNA

*shh.L* 643b*-egfp* and *shh.S* 806b*-egfp* donor plasmids were generated as previously described^S2,S3^. Genomic fragments containing the 5′ UTR of *shh.L* or *shh.S* (Figure S1B) were amplified from wild-type genomic DNA using the following primers: *shh.L* (forward: 5′-ACAGTGGCAGAGGTGTATAGAAAC-3′; reverse: 5′-CTCGTCCGAGCGAAGCCAAT-3′), *shh.S* (forward: 5′-TGTGCCTATCTCTATCTATCACAT-3′; reverse: 5′-CTCGTCCGAGCGAAGCCAAT-3′). sgRNAs targeting the 5′ UTR (Figure S1B) and coding (Figure S2A) regions of the *shh.L* and *shh.S* loci were designed using CRISPRscan^S4^ and synthesized as previously described^S2,S3^. Recombinant Cas9 protein (Alt-R S.p. Cas9 Nuclease 3NLS) was purchased from Integrated DNA Technologies (Coralville, IA, USA).

### Microinjection

Microinjection was performed as previously described^S2,S3^. Briefly, fertilized eggs were injected with 9.2 nL of injection mixture containing 800 pg of sgRNA and 2 ng of recombinant Cas9 protein in injection buffer (150 mM KCl and 20 mM HEPES, pH 7.5)^S5^. For *egfp* knock-in experiments, 50 pg of donor plasmid was included in the injection mixture. Injected embryos were maintained at 18–25°C until the desired development stages.

### Imaging

*X. laevis* embryos and larvae were anesthetized with 0.02% tricaine methanesulfonate (MS222; Sigma-Aldrich) in the rearing medium. Bright-field and fluorescence images were captured using a Nikon DS-Ri1 digital camera (Nikon, Tokyo, Japan) and a Leica MZ165FC microscope (Leica Microsystems, Wetzlar, Germany) equipped with an X-Cite XYLIS illumination system (Excelitas Technologies, Waltham, MA, USA).

### RNA extraction and qRT-PCR

Total RNA was extracted from 15 limb buds or two tails of NF stage 50–51 tadpoles using Trizol reagent (Invitrogen, Carlsbad, USA). cDNA was synthesized using the PrimeScript RT reagent Kit with gDNA Eraser (Takara) and qRT-PCR was performed using TB Green Premix Ex Taq II (Takara) on a Thermal Cycler Dice Real Time System (Takara) with the following primers: *shh.L* (forward: 5′-TAAATACGGAATGTTGGGCCG-3′, reverse: 5′-CACCAAATTCCACCATCACCCT-3′), *shh.S* (forward:5′-AAATACGGAATGTTGGCCCGA-3′, reverse: 5′-ACCAAGTTCCACCATCACCTC-3′), and *ef1α* (forward: 5′-CAGGCCAGATTGGTGCTGGATATGC-3′, reverse: 5′-GCTCTCCACGCACATTGGCTTTCCT-3′).

### Genomic DNA extraction and sequencing

Embryos or tadpoles were deeply anesthetized with 0.4% MS222 and then euthanatized for genomic DNA extraction. The whole body was lysed in a lysis buffer [10 mM Tris-HCl (pH 8.0), 5 mM EDTA, 0.5% SDS] and treated with RNase A (50 µg/mL) at 50°C for 3 h. DNA was extracted with phenol and precipitated with ethanol. Homeolog-specific sequences were amplified by PCR using PrimeSTAR Max DNA Polymerase (Takara) with the following primers: *shh.L* (forward: 5′-ACCTGCCTTGCTTGGGATCG-3′, reverse: 5′-TGCTCTCCTCGTCTTTAAACATA-3′), *shh.S* (forward: 5′-TTGTGCCCCTGTCTTGTCTG-3′, reverse: 5′-GTGTTCTCCTCGTCTTTAAATAC-3′). The PCR products were purified using DNeasy Blood & Tissue Kit (Qiagen, Hilden, Germany) and sequenced with the following primers: *shh.L* (forward: 5′-ACCTGCCTTGCTTGGGATCG-3′, reverse: 5′-TGCTCTCCTCGTCTTTAAACATA-3′), *shh.S* (forward: 5′-TTGTGCCCCTGTCTTGTCTG-3′, reverse: 5′-GTGTTCTCCTCGTCTTTAAATAC-3′). The resulting Sanger sequencing traces were analyzed using TIDE^S6^ to estimate indel frequencies relative to the corresponding wild-type sequence.

### Bone staining

Whole-mount bone-stained specimens were prepared based on a previously described rapid and nondestructive protocol for *Xenopus*^S7^, with minor modifications. Specimens were fixed overnight in 5% formalin/10% Triton X-100/1% KOH and then cleared in 10% Triton X-100/1% KOH before staining. Skeletal elements were stained with 0.05% Alizarin Red S/1% KOH and further cleared in 20% Tween 20/1% KOH before imaging.

## Acknowledgements

We are grateful to Drs. Elizabeth Nakajima and Tetsuya Nakamura for their critical reading of an earlier version of the manuscript and for providing insightful comments. This work was supported by JST SPRING, Grant Number JPMJSP2175 (to NK), Special grant for research in University of Hyogo (to MM), University of Hyogo Science and Technology Support Foundation (to MM), NIBB Collaborative Research Program (22NIBB331, 23NIBB340 to MM), the Japan Society for the Promotion of Science (JSPS), KAKENHI Grants-in-Aid for Scientific Research (B) (JP21H03829 and JP25K02288 to KTS), a Grant-in-Aid for Scientific Research on Innovative Areas (25124708 to KTS), and the Japan Science and Technology Agency (JST), CREST program (JPMJCR 2025 to KTS).

## Author contributions

Conceptualization, N.K., K.T.S. and M.M.; Investigation, N.K., R.M. and M.M.; Writing – Original Draft, N.K. and M.M.; Writing – Review & Editing, Y.U., K.T.S., and M.M.; Funding Acquisition, N.K., K.T.S. and M.M.; Supervision, Y.U. and M.M.

## Declaration of generative AI and AI-assisted technologies in the writing process

During the preparation of this manuscript, the authors used Gemini (Google) and ChatGPT (OpenAI) for grammatical correction and English language editing. After using these tools, the authors reviewed and edited the content as needed and take full responsibility for the final version of the manuscript.

## Supplemental figure legends

**Figure S1.**
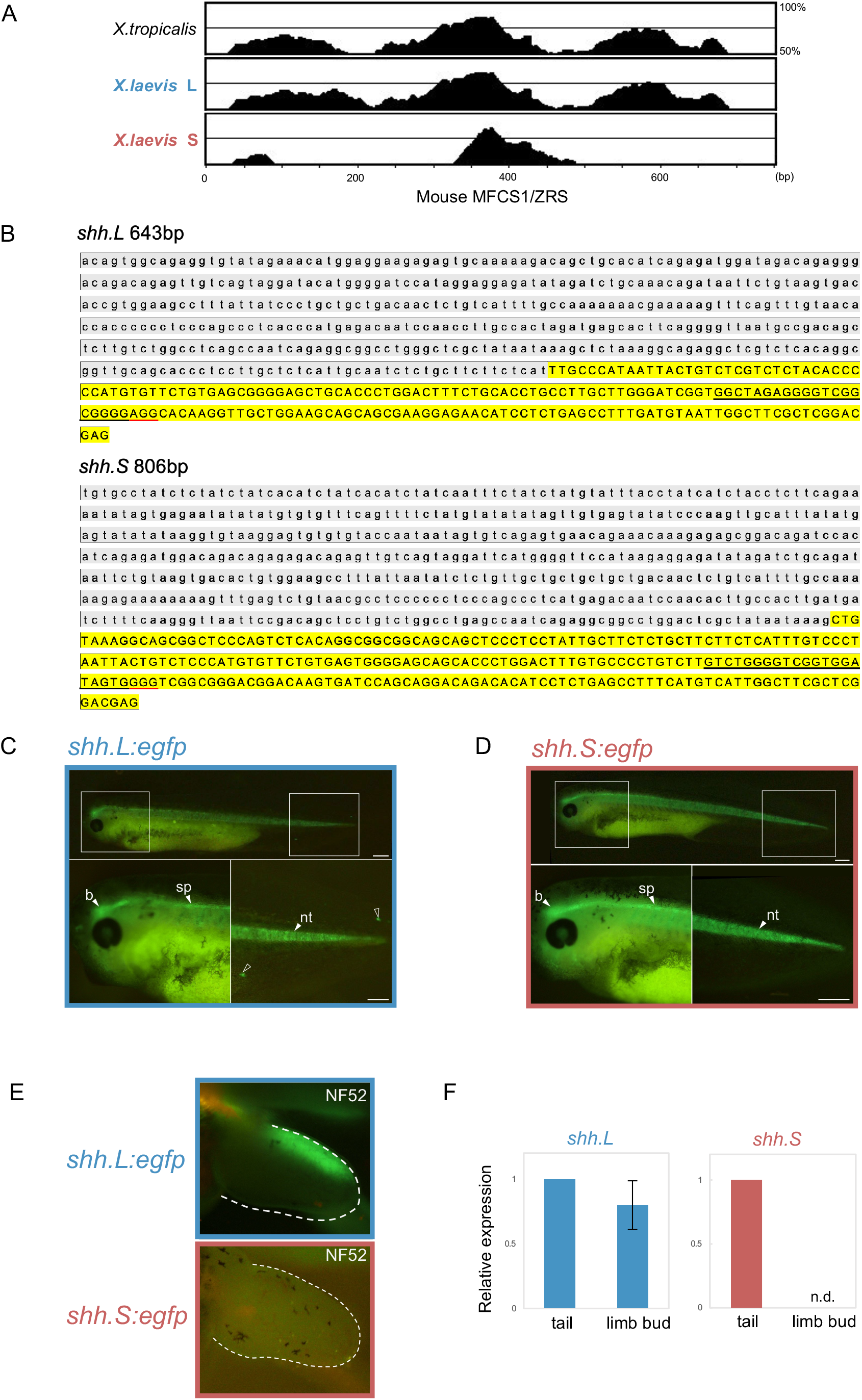
Production of *shh.L:egfp* and *shh.S:egfp Xenopus laevis* (A) VISTA alignment of MFCS1/ZRS sequences among mouse, *X. tropicalis*, and *X. laevis* L and S subgenomes. The horizontal axis represents the mouse MFCS1/ZRS core, and the vertical axis indicates sequence similarity. (B) DNA sequences of the genomic fragments included in the donor DNAs. Gray shading indicates upstream sequences, and yellow shading indicates the 5′ UTR. Black lines mark the sgRNA target sites, and red lines denote the PAM sequences. (C and D) Representative fluorescence images of *shh.L:egfp* (C) and *shh.S:egfp* (D) embryos showing specific eGFP expression at NF 37/38 in lateral views. Magnified views of the boxed regions are shown. Open arrowheads indicate ectopic eGFP signals. b, brain; nt, notochord; sp, spinal cord. (E) Representative fluorescence images of NF 52 hindlimb buds in *shh.L:egfp* (upper) and *shh.S:egfp* (lower) tadpoles. Dotted lines indicate the outlines of limb buds. Scale bar: 300 µm. (F) Relative expression of *shh.L* and *shh.S* in wild-type limb buds and tails (NF 50–51). *shh.L* expression in limb buds was detected at 0.798 relative to tail expression, whereas *shh.S* expression in limb buds was not detected (n = 3, 15 limb buds pooled per replicate). Bars represent the mean ± SD.

**Figure S2.**
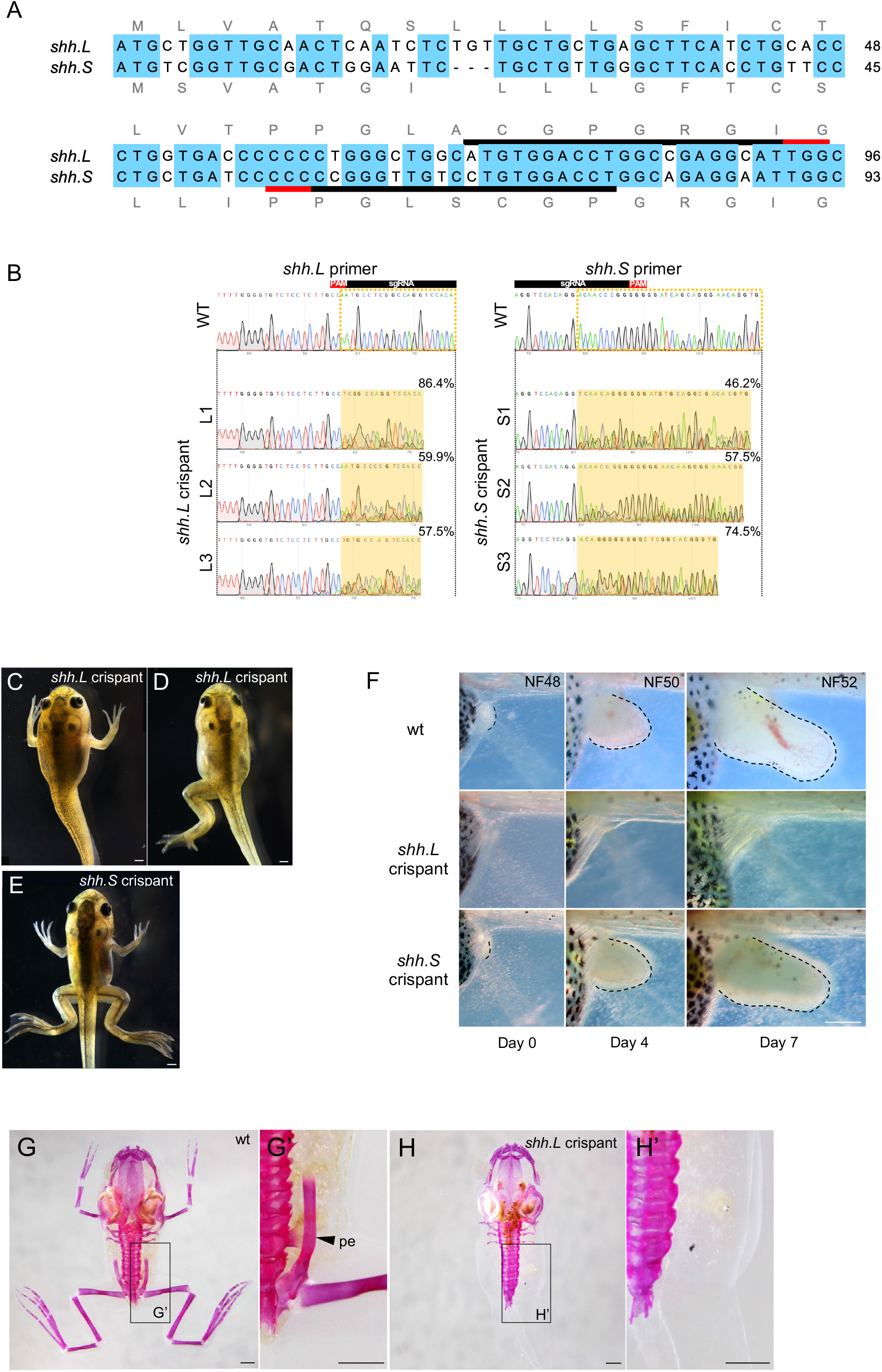
Targeted disruption of the *shh.L* or *shh.S* coding region (A) Partial genomic sequences of *shh.L* (top) and *shh.S* (bottom) downstream of the start codon. Blue shading indicates common nucleotides between *shh.L* and *shh.S*. Black bars denote the sgRNA target sequences, red bars indicate the PAM sequences. Gray letters indicate the predicted amino acid sequences. (B) Sanger sequencing chromatograms of PCR amplicons from wild-type, *shh.L* crispant, and *shh.S* crispant embryos are presented. The left panels show chromatograms and sequences spanning 20 bp upstream and downstream of the PAM in wild-type and *shh.L* crispants (L1–L3), amplified using *shh.L*-specific primers. The right panels show those of wild-type and *shh.S* crispants (S1–S3), amplified using *shh.S*-specific primers. Red and black boxes above the wild-type sequences indicate the PAM and sgRNA target sequences, respectively. Dotted and filled yellow boxes highlight regions with mixed sequencing peaks in *shh.L* and *shh.S* crispants. Percentages in the upper right indicate the total mutation frequencies estimated using Tracking of Indels by DEcomposition (TIDE)^S6^. (C) Time-course images of the hindlimb region in wild-type, *shh.L* crispant, and *shh.S* crispant. Upper panels: wild-type on Day 0 (NF 48), Day 4 (NF 50), and Day 7 (NF 52). Middle/lower panels: *shh.L* and *shh.S* crispants of comparable body lengths. Dotted lines indicate the limb buds. Scale bar: 300 µm. (D–F) Representative images of *shh.L* crispants (D and E) and an *shh.S* crispant (F) tadpole at the climax stage. (G and H) Representative bone-stained wild-type and *shh.L* crispant tadpoles at the climax stage. (G’ and H’) Magnified views of the boxed regions in (G) and (H), respectively. pe, pelvic girdle. Note that both the hindlimbs and pelvic girdles are completely absent in the *shh.L* crispant. Scale bar: 1 mm.

---

**Supplementary table 1.** eGFP expression in embryos.

| Target locus | eGFP+ / Total injected embryos | Percentage |
| --- | --- | --- |
| *shh.L* | 43/763 | 5.6% |
| *shh.S* | 36/507 | 7.1% |

Frequency of injected embryos showing specific eGFP expression in axial tissues at NF stage 40 following CRISPR-Cas9-mediated reporter targeting at the *shh.L* or *shh.S* loci.

**Supplementary table 2.** Limb bud eGFP expression in *shh.L:egfp* tadpoles.

| Target locus | Limb bud eGFP+<br>/ Reporter-positive tadpoles | Percentage |
| --- | --- | --- |
| *shh.L* | 8/35 | 22.9% |
| *shh.S* | 0/36 | 0% |

Frequency of limb bud-specific eGFP expression at NF stage 48–52 among tadpoles showing eGFP expression in axial tissues.

**Supplementary table 3.** Detailed phenotypic spectrum of limb defects in *shh.L* crispants.

| Phenotypes |  | Number of crispants (n) |
| --- | --- | --- |
| Forelimb | Hindlimb |  |
| Bilateral loss | Bilateral loss | 3* |
| Unilateral loss (Right) | Bilateral loss | 1* |
| No loss | Bilateral loss | 1* |
| Bilateral loss | No loss | 1 |
| Unilateral loss (Right) | Unilateral loss (Left) | 1 |
| Unilateral loss (Right) | No loss | 1 |
| No loss | Unilateral loss (Left) | 1 |
| No loss | No loss | 1 |
| Total (n) |  | 10 |

Individual combinations of forelimb and hindlimb phenotypes observed in *shh.L* crispants at metamorphic stage Data represent the exact count of tadpoles (n) for each respective defect combination, arranged in descending order of phenotypic severity. Asterisks indicate complete loss of pelvic girdle (n = 5).

